# *Pseudomonas aeruginosa* supports the survival of *Prevotella melaninogenica* in a cystic fibrosis lung polymicrobial community through metabolic cross-feeding

**DOI:** 10.1101/2024.10.21.619475

**Authors:** Bassam El Hafi, Fabrice Jean-Pierre, George A. O’Toole

## Abstract

Cystic fibrosis (CF) is a multi-organ genetic disorder that affects more than 100,000 individuals worldwide. Chronic respiratory infections are among the hallmark complications associated with CF lung disease, and these infections are often due to polymicrobial communities that colonize the airways of persons with CF (pwCF). Such infections are a significant cause of morbidity and mortality, with studies indicating that pwCF who are co-infected with more than one organism experience more frequent pulmonary exacerbations, leading to a faster decline in lung function. Previous work established an *in vitro* CF-relevant polymicrobial community model composed of *P. aeruginosa*, *S. aureus*, *S. sanguinis*, and *P. melaninogenica*. *P. melaninogenica* cannot survive in monoculture in this model. In this study, we leverage this model to investigate the interactions between *P. aeruginosa* and *P. melaninogenica*, allowing us to understand the mechanisms by which the two microbes interact to support the growth of *P. melaninogenica* specifically in the context of the polymicrobial community. We demonstrate a cross-feeding mechanism whereby *P. melaninogenica* metabolizes mucin into short-chain fatty acids that are in turn utilized by *P. aeruginosa* and converted into metabolites (succinate, acetate) that are cross-fed to *P. melaninogenica*, supporting the survival of this anaerobe in the CF lung-relevant model.

**Importance:** Polymicrobial interactions impact disease outcomes in pwCF who suffer from chronic respiratory infections. Previous work established a CF-relevant polymicrobial community model that allows experimental probing of these microbial interactions to achieve a better understanding of the factors that govern the mechanisms by which CF lung microbes influence each other. In this study, we investigate the interaction between *P. aeruginosa* and *P. melaninogenica*, which are two highly prevalent and abundant CF lung microbes. We uncover a cross-feeding mechanism that requires the metabolism of mucin by *P. melaninogenica* to generate short-chain fatty acids that are cross-fed to *P. aeruginosa*, and into metabolized into metabolites which are then cross-fed back to *P. melaninogenica* to support the growth of this anaerobe.

## Introduction

Cystic fibrosis (CF) is genetic disorder that affects multiple organs in the human body, including the lungs, gut, pancreas, and kidneys (1–5). CF stems from a mutation in the cystic fibrosis transmembrane conductance regulator (CFTR) gene, leading to a dysfunctional CFTR ion channel, which results in the accumulation of thickened mucus in the airway (3–5). In the lungs, the mucus accumulation can lead to the partial or complete obstruction of the airways (3, 4). In fact, more than 80% of CF-related mortality before the onset of the newest therapies was due to lung disease characterized by chronic airway infections and related inflammation (6). It has now been recognized in the literature that persons with CF (pwCF) are often colonized by multiple microorganisms concurrently, establishing seemingly unique polymicrobial communities within their lungs.

An interesting feature of the polymicrobial nature of the CF lung is that it changes over time with certain microbial species dominating the community with others becoming marginalized until rendered undetectable (7). Moreover, it has been observed that the CF polymicrobial community can be dominated by *Pseudomonas*, *Streptococcus*, and/or *Prevotella* (8–10). This finding suggests that the CF microbes influence the existence of each other *in vivo*. Therefore, understanding the mechanisms by which these organisms interact can explain the establishment and maintenance of certain members within the CF microbial community.

Previous work from our group aimed to develop an *in vitro*, CF-relevant, lung polymicrobial biofilm community model to represent the lung polymicrobial diversity among pwCF. To that end, publicly available 16S rRNA sequencing data retrieved from more than 160 clinical CF sputum samples were analyzed to identify the microbial genera that had the highest prevalence and highest abundance among pwCF and could be linked to CF disease respiratory outcomes (10, 11). The representative community is composed of *Pseudomonas aeruginosa*, *Staphylococcus aureus*, *Streptococcus sanguinis*, and *Prevotella melaninogenica* (10, 11). To translate this community to an experimental mixed-culture system, mucin-containing artificial sputum medium (ASM) (12, 13) was utilized under anoxic conditions at 37°C to grow these organisms and study their behavior in both mono- and mixed-cultures (11). The rationale behind using ASM anoxically was that the medium nutritionally mimics the CF lung environment (13) and it has been previously demonstrated that the thick mucus lining the CF lung airways creates areas of anoxia (14).

In our previous study, we noted that *P. melaninogenica* could not be recovered when grown as a monoculture in mucin-containing artificial sputum medium (ASM) under anoxic growth conditions. In contrast, *P. melaninogenica* was found to be viable when co-cultured with *P. aeruginosa*, *S. aureus*, and *S. sanguinis* as a multi-species mixed culture (11). Such a finding was also consistent with previous metabolic modeling that predicted no growth of *P. melaninogenica* as a monoculture under these culture conditions (8). The mechanism whereby *P. melaninogenica* can grow in the community, but not in monoculture, has not been explored. Here, we describe a series of experimental studies that demonstrate that metabolic cross-feeding by another member of the community, *P. aeruginosa*, allows for the survival and growth of *P. melaninogenica* in the mixed community.

## Results

### *P. aeruginosa* can support the viability of *P. melaninogenica* in a cystic fibrosis lung polymicrobial community model

Previous work from our group that aimed to develop an *in vitro*, CF-relevant, lung polymicrobial community model reported that *P. melaninogenica* could not be recovered as a monoculture in mucin-containing ASM under anoxic growth conditions. However, *P. melaninogenica* was found to be viable when cultured with *P. aeruginosa*, *S. aureus*, and *S. sanguinis* as a mixed culture (11). Such finding was also consistent with previous metabolic modeling analysis that predicted that *P. melaninogenica* should not be able to grow in monoculture in mucin-containing ASM (8). Additionally, it was predicted that *P. aeruginosa* was the only member of the community effectively cross-feeding *P. melaninogenica* (8).

To determine whether a specific member within the community was responsible for the *P. melaninogenica* growth phenotype, we co-cultured *P. melaninogenica* with the other organisms of the community in different combinations using mucin-containing ASM (11, 12) under anoxic growth conditions for 24 hours. Following incubation, the viable counts of *P. melaninogenica* were determined by serially diluting and spotting the biofilm fraction of each culture condition on the appropriate selective media, then counting the resulting colony forming units (CFUs).

We observed that *P. aeruginosa* alone was sufficient to promote the growth of *P. melaninogenica* to the same extent as the four-species mixture (∼1 × 10^6^ CFU/ml; **Figure 1A**). *P. melaninogenica* also experienced enhanced growth whenever *P. aeruginosa* was present in the mixed culture regardless of whether *S. aureus* and *S. sanguinis* were also present (**Figure S1A**). On the other hand, there were only modest (<0.5 log_10_) differences in the growth of *P. aeruginosa*, *S. aureus*, and *S. sanguinis* in the various co-culture combinations (**Figure S1B-D**).

**Figure 1.**
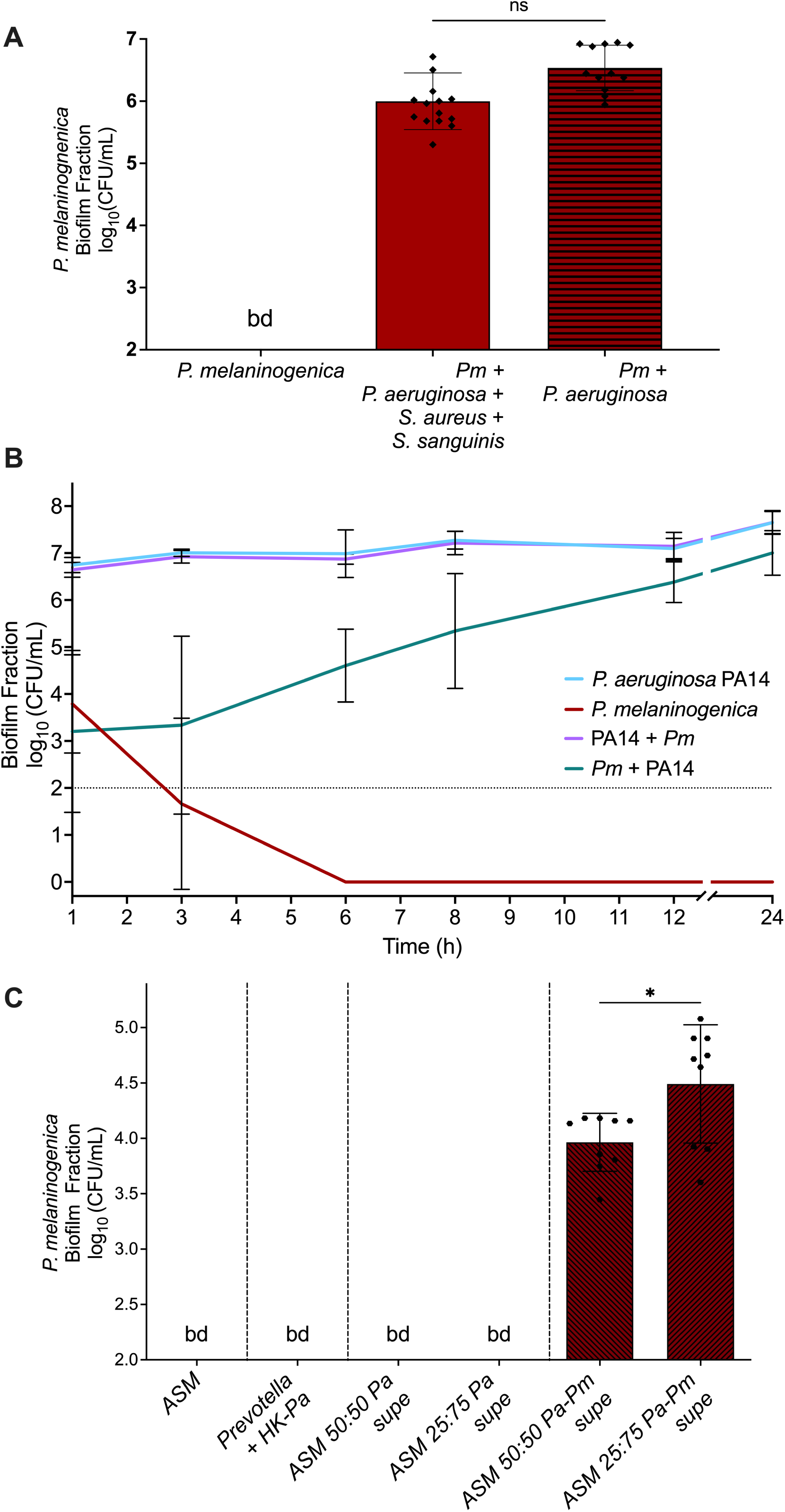
The growth of *P. melaninogenica* in co-cultures is enhanced in the presence of *P. aeruginosa*. **A**. The co-culture of *P. melaninogenica* ATCC 25845 (*Pm*) with *P. aeruginosa* PA14 (*Pa*), *S. sanguinis* SK36 (*Ss*), and *S. aureus* Newman (*Sa*). All cultures were performed using mucin-containing ASM under anoxic growth conditions at 37⁰C. The CFUs derived from the biofilm fractions of the co-cultures are plotted. Statistical significance was calculated using ordinary one-way analysis of variance (ANOVA) with Tukey’s multiple comparisons test and only one comparison is presented. **B**. A growth curve of *P. aeruginosa* PA14 and *P. melaninogenica* (*Pm*) CFUs in the biofilm fractions of their mono and co-cultures using ASM under anaerobic growth conditions at 37⁰C. The growth of *P. aeruginosa* is independent of the presence or absence of *P. melaninogenica*. The growth of *P. melaninogenica* depends on the presence of *P. aeruginosa.* Note that the growth of *P. melaninogenica* in co-culture with *P. aeruginosa* starts following a 6-hr lag period where the viable counts do not change. **C.** The monoculture of *P. melaninogenica* in mucin-containing ASM supplemented with either heat-killed *P. aeruginosa* cells or the spent ASM supernatants of *P. aeruginosa* monoculture or *P. aeruginosa*-*P. melaninogenica* co-culture at two different ratios with free ASM. All cultures were performed under anoxic growth conditions at 37⁰C. The CFUs derived from the biofilm fractions are plotted. Statistical significance was calculated using ordinary one-way analysis of variance (ANOVA) with Tukey’s multiple comparisons test and only one comparison is presented, * p < 0.05.

To understand the dynamics of the interaction between *P. aeruginosa* and *P. melaninogenica*, we conducted a time course assay of the monocultures and co-culture of both organisms in mucin-containing ASM under anoxic growth conditions, recording the CFU/mL of the biofilm fractions of the cultures at the 1-, 3-, 6-, 8-, 12-, and 24-hour time points. We observed that *P. aeruginosa* was not affected by the presence of *P. melaninogenica* since the growth curve of *P. aeruginosa* was nearly identical in both mono- and co-culture conditions, reaching ∼10^8^ CFU/mL by the 24-hr time point (**Figure 1B**). In contrast, *P. melaninogenica* was not able to survive the monoculture condition as its viable population decreased over time until it was no longer detectable at the 6-hr time point. However, in co-culture with *P. aeruginosa*, *P. melaninogenica* appeared to require a 6-hour adjustment or lag period before steadily growing over the next 18 hours to ∼10^6^ CFU/ml (**Figure 1B**).

The interaction between *P. aeruginosa* and *P. melaninogenica* was also not restricted to the biofilm fraction of the co-culture since the growth curves of *P. aeruginosa* and *P. melaninogenica* in the planktonic fraction of both mono- and co-culture conditions mirrored those in the biofilm fraction (**Figure S1E**).

To determine whether *P. aeruginosa* was promoting the growth of *P. melaninogenica* through secreted products, we grew *P. melaninogenica* as a monoculture in mucin-containing ASM with either heat-killed *P. aeruginosa* cells or different ratios of spent *P. aeruginosa* monoculture ASM supernatant as well as spent *P. aeruginosa*-*P. melaninogenica* co-culture ASM supernatant. We observed that *P. melaninogenica* only survived monoculture when supplemented with the spent *P. aeruginosa*-*P. melaninogenica* co-culture ASM supernatant **(Figure 1C)**. These data imply that, firstly, *P. aeruginosa* supports the growth of *P. melaninogenica* through a mechanism that requires live cells to secrete sharable products given that heat-killed cells could not rescue *P. melaninogenica* growth. Secondly, these secreted products can only rescue *P. melaninogenica* when both organisms are in co-culture in mucin-containing ASM - the spent *P. aeruginosa* monoculture supernatants could not support *P. melaninogenica* survival in monoculture.

Together, these observations demonstrate the dynamic nature of the interaction between these organisms and support a model whereby *P. melaninogenica* benefits from the presence of *P. aeruginosa* in co-culture via shared excreted products.

### A genetic screen identifies *P. aeruginosa* mutants unable to fully support *P. melaninogenica* growth when co-cultured in mucin-containing ASM

We next sought to investigate the mechanisms that govern the interaction between *P. aeruginosa* and *P. melaninogenica* using a genetic approach. To achieve this, we screened the *P. aeruginosa* PA14 non-redundant transposon mutant library (15) in co-culture with *P. melaninogenica,* using mucin-containing ASM under anoxic conditions, to identify *P. aeruginosa* PA14 transposon mutants that were incapable of, or showed a reduced ability for, supporting the growth of *P. melaninogenica* (**Figure 2A**). We reasoned that this approach would allow us to identify genetic determinants of *P. aeruginosa* that are required for its ability to promote the growth of *P. melaninogenica* in co-culture.

**Figure 2.**
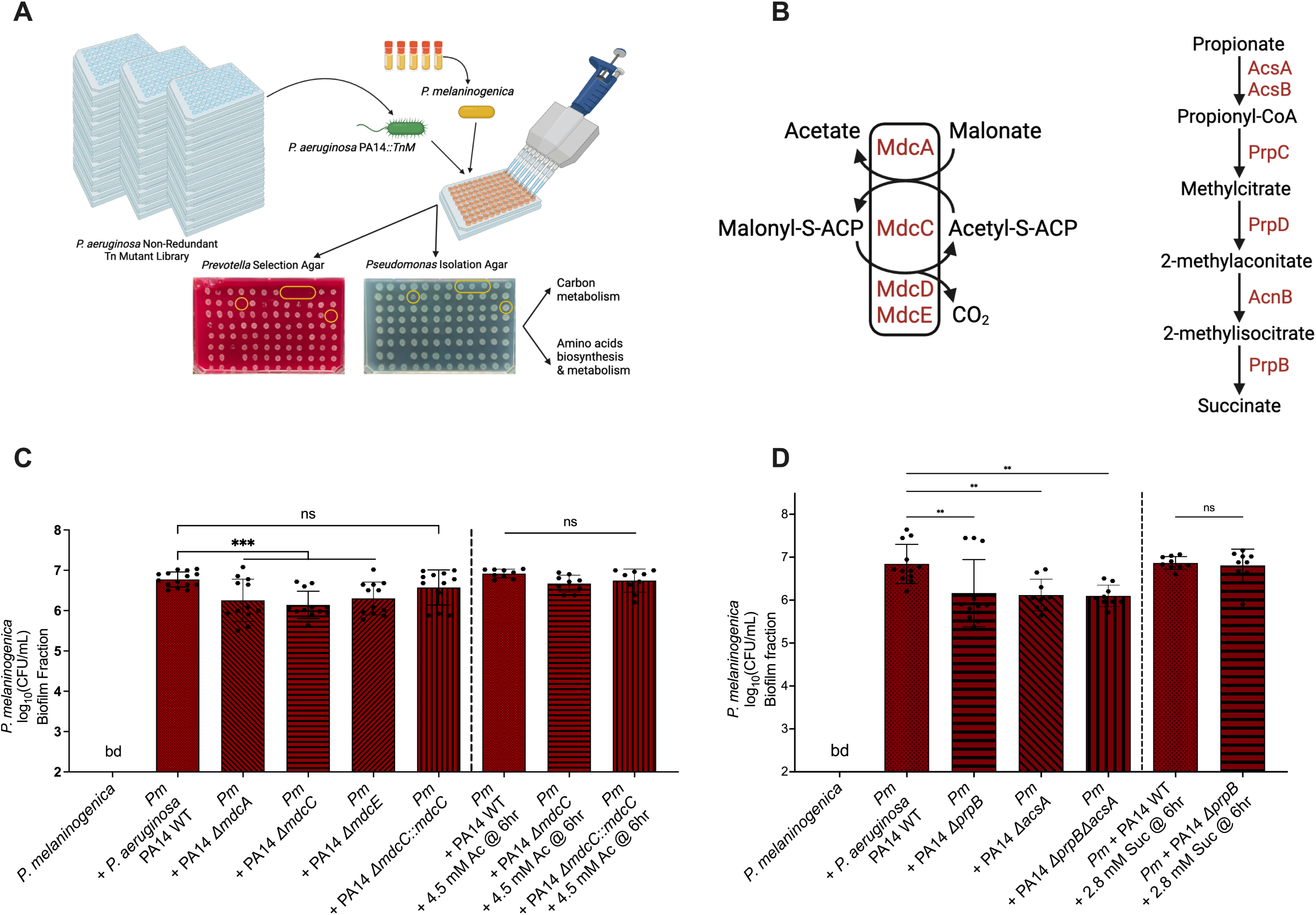
*P. aeruginosa* genetic screen identifies carbon metabolism pathways as being implicated in the interaction with *P. melaninogenica*. **A.** A schematic diagram of the *P. aeruginosa* PA14 non-redundant transposon mutant library screen in co-culture with *P. melaninogenica*. Figure designed using BioRender. **B.** Pathways showing the metabolism of malonate (21, 52) and propionate (21) into acetate and succinate, respectively, with selected genes involved in the catabolism of these carbon sources. **C-D.** Bar plots of the CFUs derived from the biofilm fractions of *P. melaninogenica*. All cultures were performed using mucin-containing ASM under anaerobic growth conditions at 37⁰C. Statistical significance was calculated using ordinary one-way analysis of variance (ANOVA) with Tukey’s multiple comparisons test. **C**. The pairwise co-culture of *P. melaninogenica* with WT *P. aeruginosa* PA14 and the Δ*mdcA*, Δ*mdcE,* and Δ*mdcC* mutants as well as the Δ*mdcC*::*mdcC* complemented strain, with and without the supplementation of 4.5 mM of acetate at 6 hrs, *** p < 0.05. The concentrations of acetate used here was selected based on the data presented in Figure 4. **D**. The pairwise co-culture of *P. melaninogenica* with WT *P. aeruginosa* PA14 and Δ*prpB*, Δ*acsA*, and Δ*prpB*Δ*acsA* mutants, with and without the supplementation of 2.8 mM of succinate at 6 hrs, ** p < 0.05. The concentrations of succinate used here was selected based on the data presented in Figure 4.

Mutant candidates that were identified in the primary screen were subjected to a second and third round of re-testing to generate the final list of *P. aeruginosa* PA14 transposon mutants that showed reduced ability to support the growth of *P. melaninogenica* in co-culture compared to the wild-type (WT) *P. aeruginosa* PA14 in mucin-containing ASM during anaerobic growth (**Table S1**). Many of the identified *P. aeruginosa* PA14 mutants from our screen were genes of unknown function. However, a few mutations were in genes with defined functions, such as *sppR*, which encodes the TonB-dependent receptor SppR. The *sppR* gene is co-transcribed with the *spp* operon, which is responsible for the expression of the Spp transporter, involved in xenosiderophore uptake (16, 17). Another mutation mapped to the *cupD4* gene, which encodes an adhesin in the CupD fimbrial assembly (16, 18).

A number of mutations were in genes involved in either carbon metabolism or amino acid biosynthesis and metabolism, namely: (i) *mdcE*, which encodes a subunit of malonate decarboxylase (16, 19–21), (ii) *prpR*, which encodes the transcriptional activator of the *prp* genes that encode the enzymes responsible for metabolizing propionate to succinate (16, 21), (iii) *hutU*, which is part of the histidine utilization locus and encodes urocanate hydratase that converts urocanate into imidazolone propionate as part of the histidine catabolism pathway by *Pseudomonas fluorescens* (16, 22), and (iv) PA14_38140, which is the ortholog of the *pauA4* gene of *P. aeruginosa* PAO1 (16) that encodes glutamylpolyamine synthetase involved in polyamine metabolism (23).

Identifying genes in our genetic screen with defects in these metabolic pathways is consistent with metabolic modeling studies that were previously conducted by our group (8, 11). Those modeling efforts identified several organic acids and amino acids that were predicted to be cross-fed between the members of the CF-relevant polymicrobial community. Therefore, we decided to focus on select *P. aeruginosa* PA14 metabolism-related pathways identified by the genetic screen to better understand their role in the interaction between *P. aeruginosa* and *P. melaninogenica*.

### Acetate and succinate contribute to the growth of *P. melaninogenica* when co-cultured with *P. aeruginosa* in mucin-containing ASM

In *P. aeruginosa*, malonate is decarboxylated to acetate via malonate decarboxylase (**Figure 2B**, left), which is composed of multiple subunits encoded by the *mdcABCDEGHLM* operon. The *mdcABCDEGH* genes encode functional subunits of the Mdc enzyme complex and the *mdcLM* genes encode the malonate transporter (20). Propionate, on the other hand, is metabolized via the methylcitrate pathway (**Figure 2B**, right), which is composed of a number of intermediary steps, starting with the conversion of propionate to propionyl-CoA via acetyl-CoA synthetase, encoded by *acsA*, and ending with the conversation of 2-methylisocitrate into succinate via 2-methylisocitrate lyase, encoded by the *prpB* gene (16, 21, 24–26).

To examine the involvement of malonate and propionate metabolism pathways in the interaction between *P. aeruginosa* and *P. melaninogenica*, we acquired previously reported *P. aeruginosa* PA14 strains carrying deletion mutations of three different *mdc* genes that encode the active site subunits of Mdc (20), as well as the deletion mutants of the *acsA* and *prpB* genes, which respectively encode AcsA and PrpB, required for the metabolism of propionate to succinate (27). We co-cultured the selected mutants with *P. melaninogenica* in mucin-containing ASM under anoxic growth conditions, then counted the resulting CFUs of each organism on selective media and compared those results to the growth of *P. melaninogenica* in co-culture with WT *P. aeruginosa* PA14.

The co-culture of *P. melaninogenica* with the *P. aeruginosa* PA14 Δ*mdcA,* Δ*mdcC* and Δ*mdcE* mutants resulted in a significant, ∼10-fold decrease in viable *P. melaninogenica* counts when compared to its co-culture with WT *P. aeruginosa* PA14 (**Figure 2C**). Additionally, co-culturing *P. melaninogenica* with the *P. aeruginosa* PA14 Δ*mdcC::mdcC,* a complemented strain, restored the viable counts of *P. melaninogenica* to a level not significantly different from WT *P. aeruginosa* PA14 (**Figure 2C**).

We then asked whether supplementing the end-product of malonate catabolism (i.e., acetate) would reverse the growth defect of *P. melaninogenica* when co-cultured with the *mdc* mutants. Interestingly, adding 4.5 mM of acetate to the *P. melaninogenica*-*P. aeruginosa* PA14 Δ*mdcC* co-culture at the 6-hr time point, but not at the start of the experiment (t=0, not shown), restored the growth of *P. melaninogenica* to levels observed in co-culture with WT *P. aeruginosa* PA14 (**Figure 2C**). The 6-hr time point was selected because, as we noted earlier, *P. melaninogenica* growth in co-culture with WT *P. aeruginosa* PA14 appeared to increase starting at the 6-hr time point in the time course assay (**Figure 1B**). We address this observation further in the Discussion.

Similar to our observations analyzing the *mdc* mutants, the co-culture of *P. melaninogenica* with *P. aeruginosa* PA14 Δ*prpB*, Δ*acsA*, and Δ*prpB*Δ*acsA* resulted in a significant, ∼10-fold decrease in the viable counts of *P. melaninogenica* compared to its growth with WT *P. aeruginosa* PA14 (**Figure 2D**). Moreover, the supplementation of the *P. aeruginosa* PA14 Δ*prpB*-*P. melaninogenica* co-culture with 2.8 mM succinate, the metabolic end-product of propionate, at the 6-hr timepoint, also restored the growth of *P. melaninogenica* to levels observed in co-culture with WT *P. aeruginosa* PA14 (**Figure 2D**). The growth of wildtype *P. aeruginosa* PA14 as well the metabolic mutants was not substantially different under any of these conditions (**Figure S2A-B**).

We next tested whether the addition of 4.5 mM acetate and 2.8 mM succinate, separately and as a cocktail, to mucin-containing ASM would be sufficient to rescue *P. melaninogenica* monocultures; however, *P. melaninogenica* remained unrecoverable in monoculture after 24 hours of anoxic incubation regardless of the metabolite supplementation (**Figure S2C**).

To investigate the impact of disrupting additional metabolic pathways responsible for generating acetate and succinate in *P. aeruginosa* PA14 (**Figure 3A**), we created single and combination deletion mutants in the Δ*mdcC* and Δ*prpB* mutant backgrounds. The genes deleted include *pauA*, which encodes pimeloyl-CoA synthetase, which is involved in converting acetyl-CoA to acetate, the *sucDC* operon, which encodes succinyl-CoA synthetase that metabolizes succinyl-CoA to succinate, and the *sdhBADC* operon that encodes succinate dehydrogenase that converts fumarate to succinate (24–26). In total, nine additional deletion mutants were generated: *P. aeruginosa* PA14 Δ*pauA*, Δ*sucDC*, Δ*prpB*Δ*mdcC*, Δ*sucDC*Δ*prpB*, Δ*sucDC*Δ*mdcC*, Δ*mdcC*Δ*pauA*, Δ*sucDC*Δ*prpB*Δ*mdcC*, Δ*prpB*Δ*mdcC*Δ*pauA*, and Δ*prpB*Δ*mdcC*Δ*sdhBADC*.

**Figure 3.**
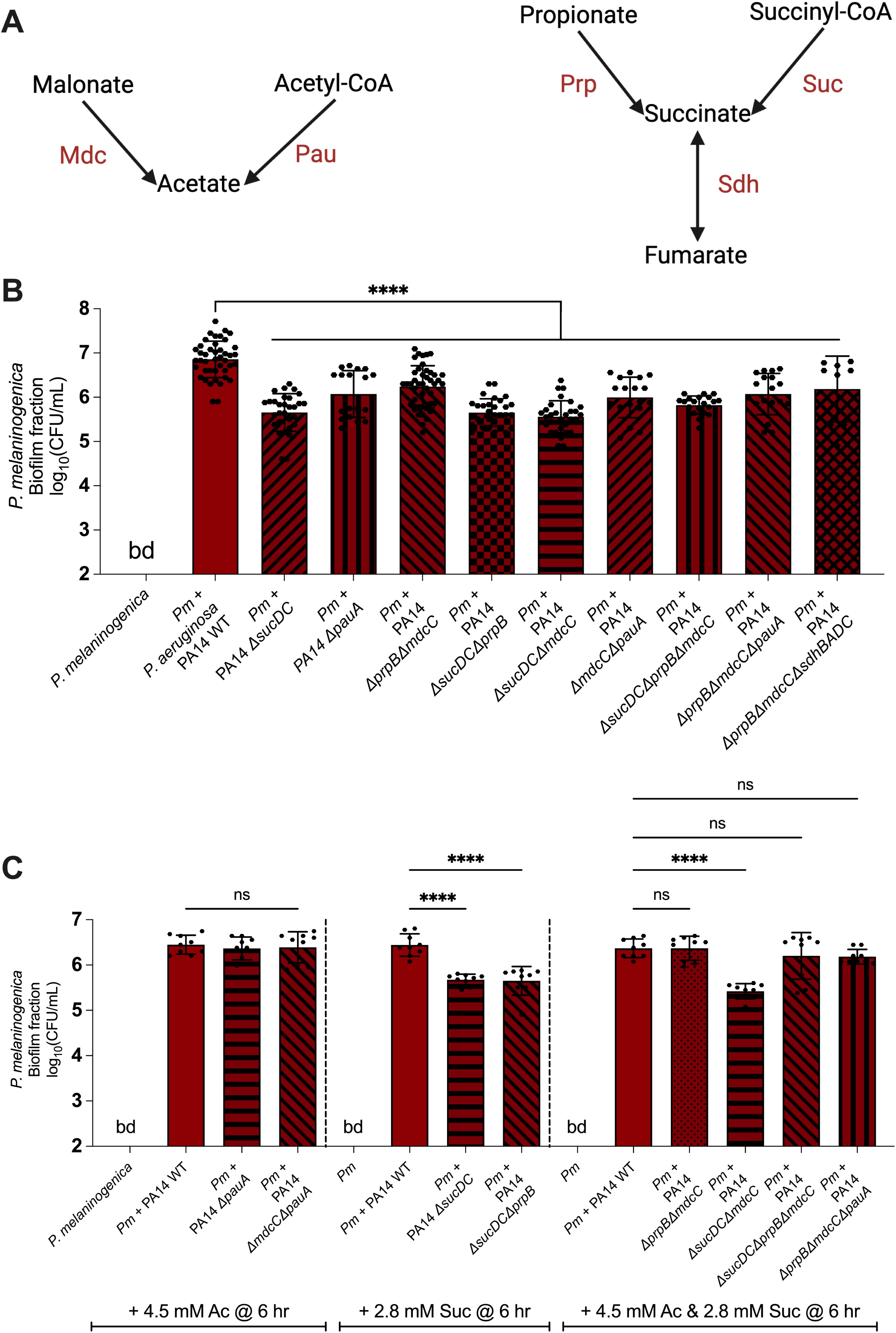
Additional metabolic pathways that generate acetate and succinate in *P. aeruginosa* are also implicated in the interaction with *P. melaninogenica*. **A.** A schematic diagram indicating multiple metabolic pathways that generate acetate and succinate in *P. aeruginosa*. **B-C**. Bar plots of the CFUs derived from the biofilm fractions of *P. melaninogenica*. All cultures were performed using mucin-containing ASM under anaerobic growth conditions at 37⁰C. Statistical significance was calculated using ordinary one-way analysis of variance (ANOVA) with Tukey’s multiple comparisons test. **B.** The co-culture of *P. melaninogenica* with WT *P. aeruginosa* PA14 or the Δ*sucDC*, Δ*pauA*, Δ*prpB*Δ*mdcC*, Δ*sucDC*Δ*prpB,* Δ*sucDC*Δ*mdcC*, Δ*mdcC*Δ*pauA*, Δ*sucDC*Δ*prpB*Δ*mdcC*, Δ*sucDC*Δ*prpB*Δ*pauA*, and Δ*sucDC*Δ*prpB*Δ*sdhBADC* mutants, **** p < 0.0001. **C**. Selected strains from panel B supplemented, as indicated, with acetate (left), succinate (middle) or both (right), **** p < 0.0001.

Upon co-culturing *P. melaninogenica* with the additional PA14 mutants in mucin-containing ASM under anoxic growth conditions, we noted a significant, ∼10-15-fold decrease in the recovery of *P. melaninogenica* compared to its co-culture with WT *P. aeruginosa* PA14 at the 24-hr time point (**Figure 3B**), with the largest effect being observed when the *sucDC* genes are deleted either as a single or combination mutant. Moreover, upon supplementing the co-cultures with acetate and/or succinate at the 6-hr time point, depending on whether each or both metabolic pathways were disrupted, we observed that the growth of *P. melaninogenica* was fully rescued except in the co-cultures with PA14 Δ*sucDC*, Δ*sucDC*Δ*prpB*, and Δ*sucDC*Δ*mdcC*, which still showed a growth defect in co-culture (**Figure 3C**). Lastly, it is important to note that the *P. aeruginosa* propionate and malonate metabolism mutants used here showed no significant changes in viable counts when grown in mono- or co-cultures (**Figures S3**).

### Acetate and succinate are elevated in the *P. aeruginosa*-*P. melaninogenica* co-culture compared to monocultures

Given that mutations in the pathways required for the metabolism of malonate and propionate by *P. aeruginosa* alters its interaction with *P. melaninogenica*, we sought to measure the concentration of these metabolites, as well as their respective products generated by the Mdc and Prp pathways, acetate and succinate. We quantified the endpoint concentrations these metabolites in the cell-free supernatants of the mono- and co-cultures of *P. aeruginosa* and/or *P. melaninogenica* in mucin-containing ASM by GC-MS. We observed that malonate was undetectable in all culture conditions, and propionate was in the low micromolar range, in both mono- and co-culture supernatants (**Figure 4**). The concentration of acetate was 1.05 mM in the *P. aeruginosa* PA14 monoculture supernatant and 0.06 mM in the *P. melaninogenica* monoculture supernatant; however, acetate was significantly higher in the co-culture supernatant at 4.5 mM (**Figure 4**). Succinate was found to be in the low micromolar concentration range for both monoculture supernatants but was significantly higher in the co-culture supernatant at 2.8 mM (**Figure 4**). The detection of higher acetate and succinate concentrations in the co-culture conditions of our *in vitro* model appears to be physiologically relevant in the context of CF lung infections as we address further in the Discussion.

**Figure 4.**
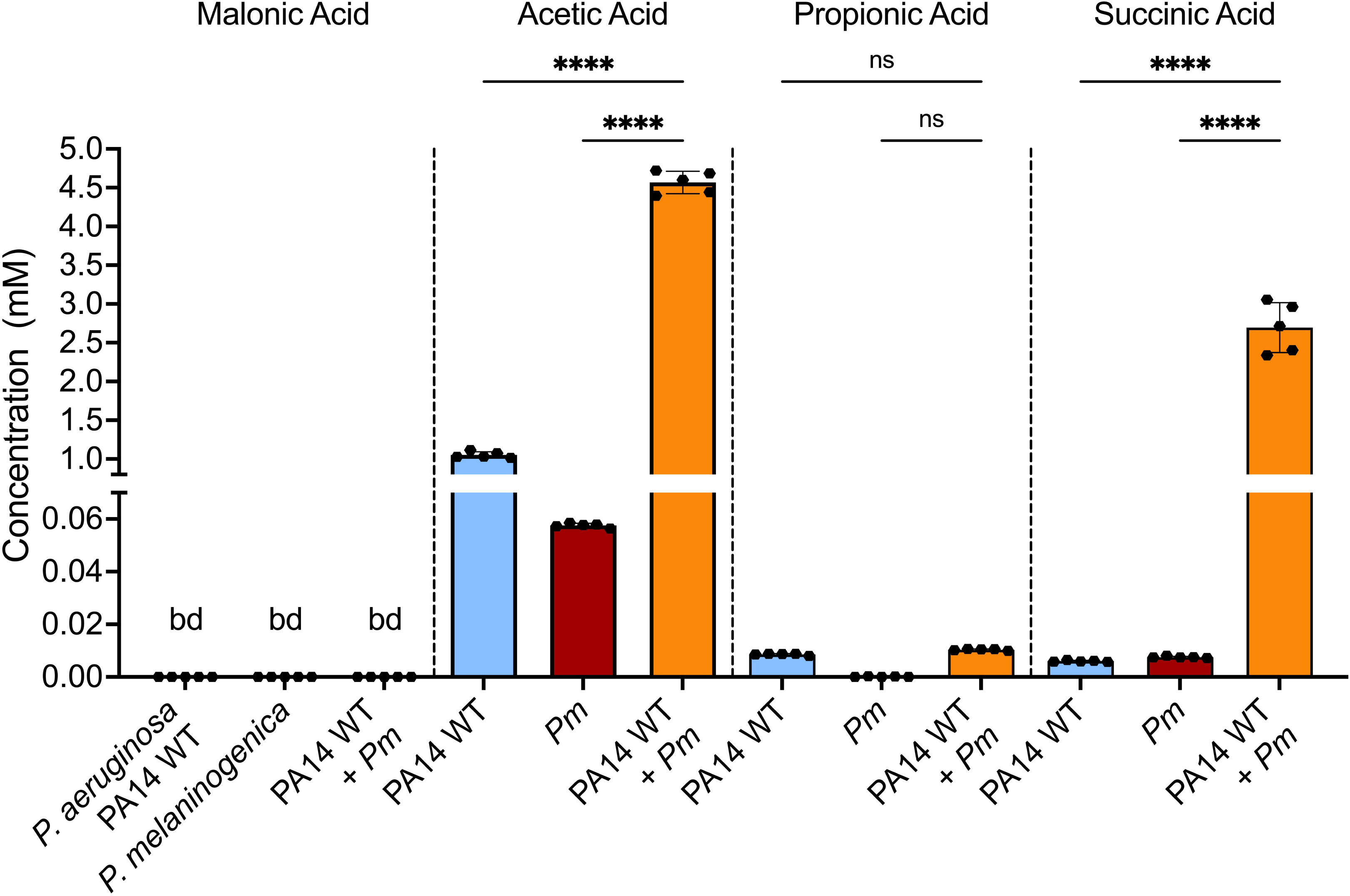
Acetate and succinate concentrations are significantly higher in *P. aeruginosa-P. melaninogenica* co-cultures compared to monoculture growth. The concentrations of malonate, acetate, propionate, and succinate as measured by GC-MS in the cell-free mono and co-culture ASM supernatants of WT *P. aeruginosa* PA14 and *P. melaninogenica* at the 24-hour time point. Statistical significance was calculated using ordinary one-way analysis of variance (ANOVA) with Tukey’s multiple comparisons test, **** p < 0.0001.

Since ASM is a complex medium composed of amino acids, sugars, mucin, metal ions, and DNA (12) that can complicate the detection of metabolites in the cell-free supernatants, we sought to simplify the co-culture medium by replacing mucin-containing ASM with M63 minimal salts base medium, supplemented with 0.2% glycerol as an energy source to support *P. aeruginosa* growth, 100 μM nitrate as the anaerobic terminal electron acceptor for *P. aeruginosa*, and mucin at the same concentration used in ASM. The viability of *P. aeruginosa* and *P. melaninogenica* mono- and co-culture was assayed by recording the biofilm fraction, and as shown in **Figure S4A**, this medium replicated the phenotype of mucin-supplemented ASM, with *P. melaninogenica* only growing in co-culture.

Using this simplified medium, we followed the concentration of the detected metabolites over time using high-pressure ion chromatography (HPIC). First, propionate was below the limit of detection in all culture conditions (**Figure S4B-D**). Secondly, the concentration of malonate appeared to increase during the early phase to approximately 0.05 mM then gradually decline as time elapsed in all culture conditions (**Figure S4B-D**). Conversely, the concentration of succinate remained low in all conditions at the early stages before climbing to 0.1 mM in the *P. melaninogenica* monoculture and approximately 0.2 mM in the co-culture; however, it remained at or near the limit of detection in the *P. aeruginosa* monoculture (**Figure S4B-D**). Finally, the concentration of acetate fluctuated around 0.6 mM in the *P. melaninogenica* monoculture but appeared to increase over time in both the *P. aeruginosa* monoculture, to reach 0.85 mM, and the co-culture to reach 1.6 mM at the 24-hr time point (**Figure S4B-D**). Thus, in both media, succinate and acetate accumulated to the highest levels in co-culture conditions.

### Carbon catabolite repression contributes to the *P. melaninogenica*-*P. aeruginosa* interaction through C_4_-dicarboxylate transport

To assess the effect of disrupting global metabolic pathways on the interaction between *P. aeruginosa* and *P. melaninogenica*, we performed co-culture experiments using *P. aeruginosa* PA14 carbon catabolite repression (CCR) mutants. In *P. aeruginosa*, CCR is a post-transcriptional metabolic regulatory process that establishes a hierarchy of preference towards the consumption of carbon sources (28). The two-component signaling system CbrAB, as well as the catabolite repression control protein Crc, are critical in the *P. aeruginosa* CCR system (28). Therefore, we acquired previously-reported *P. aeruginosa* PA14 Δ*cbrA,* Δ*cbrB* and Δ*crc* mutants (29), and co-cultured them with *P. melaninogenica*. There was a modest, but statistically significant, decrease in the growth of *P. melaninogenica* when co-cultured with the *P. aeruginosa* PA14 CCR mutants compared to WT (**Figure S5A**). None of these mutants had a growth defect in monoculture at the same 24-hr time point (**Figure S5B**). Overall, defects in the CCR system of *P. aeruginosa* alters its interaction with *P. melaninogenica*.

We additionally investigated the involvement of a variety CCR targets (30) in the interaction between *P. aeruginosa* and *P. melaninogenica* by co-culturing this anaerobe with *P. aeruginosa* PA14 *dctA::Tn*M, Δ*phzA1*Δ*phzA2*, Δ*mvfR*/*pqsR*, and Δ*pqsH* mutants. Only the mutation in strain with a mutation in *dctA*, which encodes a C_4_-dicarboxylate transporter of fumarate, malate, and succinate (31), showed a significant reduction in *P. melaninogenica* viability compared to WT PA14 (**Figure S6**).

### Mucin cannot support the growth of *P. aeruginosa* in our experimental system, consistent with previous findings

While we were able to measure the concentrations of acetate and succinate in the cell-free supernatants of our *in vitro* model as described in the previous section, the source of those compounds, as well as their respective malonate and propionate precursors, was still in question since they are not components of the culture medium we prepared. Therefore, we first hypothesized that perhaps mucin was metabolized to generate malonate and propionate, which was in turn utilized by *P. aeruginosa*.

Previous work by Flynn *et al*. (27) showed that *P. aeruginosa* cannot utilize mucin as a sole carbon source. We observed similar findings here by culturing WT *P. aeruginosa* PA14 anaerobically in minimal medium with mucin as the sole carbon/energy source and nitrate as the electron acceptor, and compared its biofilm fraction CFU counts after 24 hours to those recorded from the culture in minimal media lacking mucin, as well as to mucin-containing ASM as a positive control. We found that mucin, as a sole energy source, did not significantly impact the growth of *P. aeruginosa* in the minimal medium since it maintained its initial inoculum of ∼10^6^-10^7^ CFU/mL (dotted line) after 24 hours without significantly increasing or decreasing its viability, whether or not mucin was present (**Figure 5A**). These results indicate that *P. aeruginosa* does not effectively utilize mucin as a carbon source; thus, was unlikely able to generate the metabolic intermediates (i.e., malonate and propionate) implicated in its interaction with *P. melaninogenica*.

**Figure 5.**
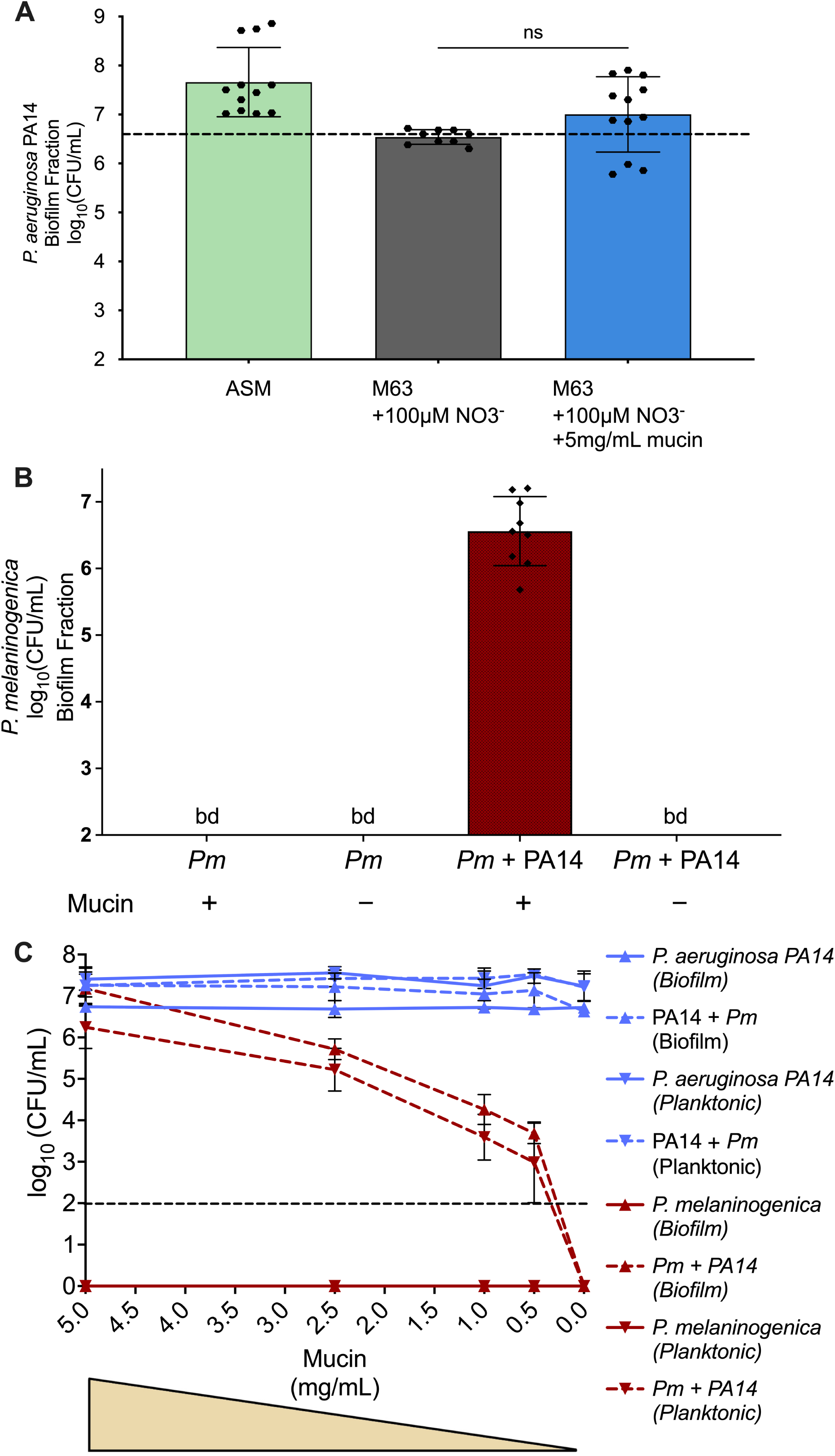
Mucin is required for *P. aeruginosa* promotion *of P. melaninogenica* growth. **A**. The monoculture of *P. aeruginosa* PA14 in ASM compared to minimal medium supplemented with nitrate +/- mucin as the main carbon source conducted under anaerobic growth conditions at 37⁰C. The dashed line indicates the starting inoculum concentration of *P. aeruginosa*. The CFUs of the biofilm fraction of *P. aeruginosa* PA14 are plotted. Statistical significance was calculated using ordinary one-way analysis of variance (ANOVA) with Tukey’s multiple comparisons test. No statistical significance was found. **B**. The mono- and co-culture of *P. melaninogenica* with *P. aeruginosa* PA14 in the presence and absence of mucin in ASM. The CFUs of the biofilm fractions of *P. melaninogenica* are plotted from an experiment conducted under anaerobic growth conditions at 37⁰C. *P. aeruginosa* does not support the growth of *P. melaninogenica* in the absence of mucin. **C**. The mono and co-cultures of *P. melaninogenica* and *P. aeruginosa* PA14 in ASM with decreasing concentrations of mucin. The CFUs of both planktonic and biofilm fractions are plotted. The experiment was conducted under anaerobic growth conditions at 37⁰C. The survival of *P. melaninogenica* in co-culture with *P. aeruginosa* depends on mucin. The dashed line indicates the limit of detection.

### Mucin is a critical factor mediating the *P. aeruginosa*-*P. melaninogenica* interaction under CF-like nutritional environments

To assess whether mucin is required for *P. aeruginosa* to support the growth of *P. melaninogenica* in our *in vitro* model, we grew the organisms in mono- and co-cultures in ASM with and without mucin, then enumerated the resulting biofilm fraction CFUs after 24 hours of anoxic incubation at 37°C. As expected, *P. melaninogenica* was not detectible when grown as a monoculture +/- mucin (**Figure 5B**). By contrast, *P. melaninogenica* was recoverable at approximately 4×10^6^ CFU/mL when co-cultured with *P. aeruginosa* in mucin-containing ASM (**Figure 5B**). However, *P. melaninogenica* growth was no longer observed in co-culture with *P. aeruginosa* when ASM lacking mucin was used (**Figure 5B**). This observation demonstrated the necessity of mucin for the survival of *P. melaninogenica* in co-culture with *P. aeruginosa*.

*P. aeruginosa* PA14 was recoverable in the biofilm fraction of both monocultures and co-cultures, with and without mucin, at ∼10^7^ CFU/mL. In monoculture, there was no significant difference in CFU +/- mucin, while in co-culture there was a significant but modest increase (<0.5 log_10_) in *P. aeruginosa* PA14 viability with mucin (**Figure S7**).

To further investigate the dependence of *P. melaninogenica* on mucin in the co-culture medium, we anoxically grew the mono- and co-cultures of *P. aeruginosa* and *P. melaninogenica* in ASM with decreasing concentrations of mucin (5, 2.5, 1, 0.5 and 0 mg/mL), then plotted the resulting biofilm and planktonic CFUs at the 24-hr timepoint. It was evident that the growth of WT *P. aeruginosa* PA14 was neither affected by the concentration of mucin, nor by the presence or absence of *P. melaninogenica* (**Figure 5C**). Unsurprisingly, *P. melaninogenica* alone did not survive the monocultures in either biofilm or planktonic fractions regardless of the concentration of mucin (**Figure 5C**). However, *P. melaninogenica* appeared to exhibit a dose-dependent response to the concentration of mucin in both biofilm and planktonic fractions of the co-culture, where the decrease in the concentration of mucin directly correlated with the decrease in the viability of *P. melaninogenica* in co-culture with *P. aeruginosa* from ∼10^7^ to 0 CFU/mL (**Figure 5C**). Together, these observations further supported the reliance of *P. melaninogenica* on mucin for survival and growth in co-culture with *P. aeruginosa*.

### *P. melaninogenica* expresses genes implicated in mucin catabolism when grown with *P. aeruginosa* in mucin-containing ASM

Previous work from our lab investigated the transcriptional profiles of *P. aeruginosa*, *S. aureus*, *S. sanguinis*, and *P. melaninogenica* as part of the CF-relevant polymicrobial community in ASM with and without mucin (32). Upon visualizing the differential expression data of *P. melaninogenica* when grown as part of the community, it was evident that the presence of mucin was a key factor leading to the differential expression of multiple genes implicated in cellular metabolism (**Figure 6A**).

**Figure 6.**
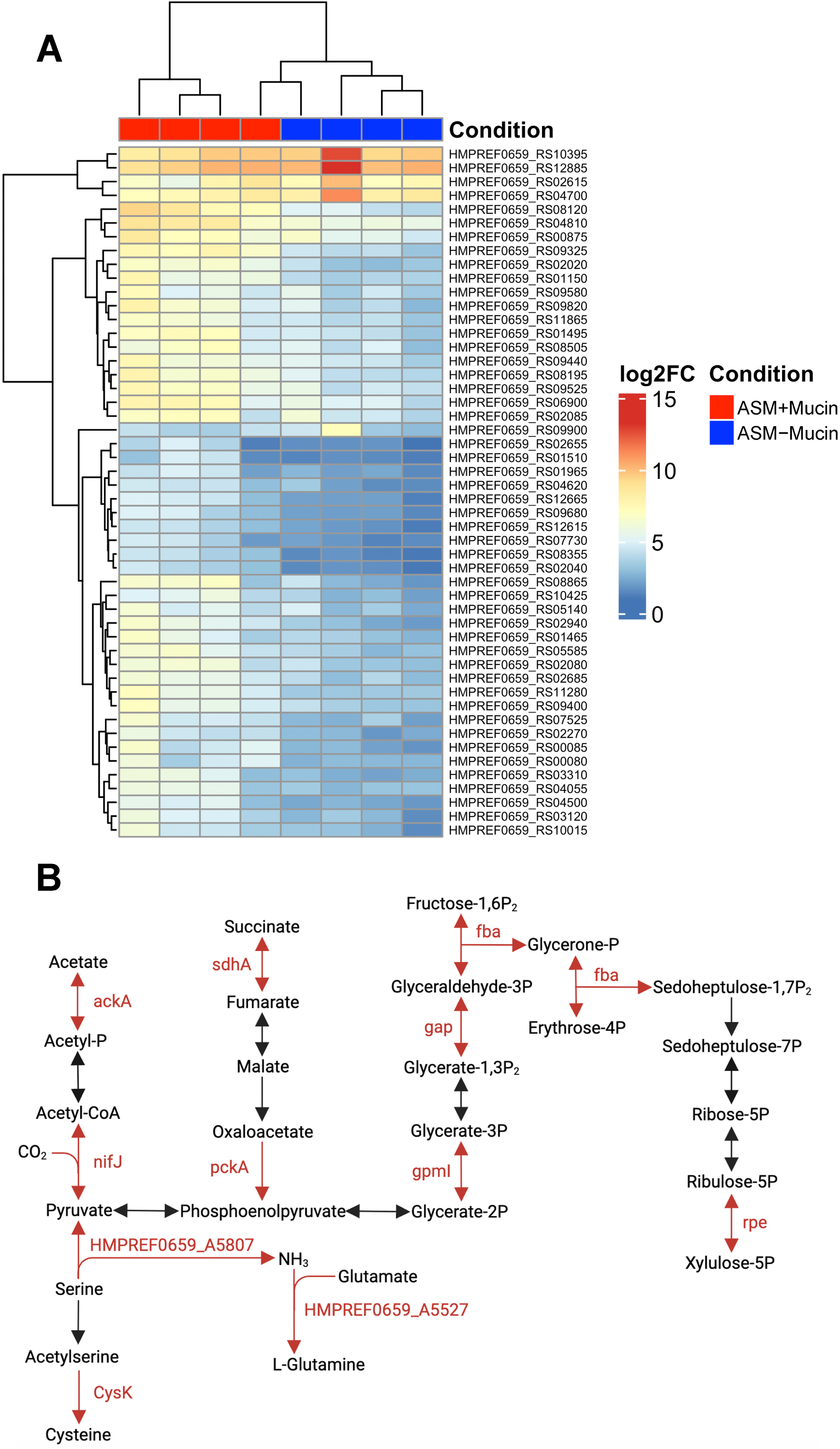
*P. melaninogenica* differentially responds to mucin in the culture medium when co-cultured the CF polymicrobial community. **A-B.** The RNAseq data used to generate the figures were originally reported in Kesthely *et al*. (32) **A.** Heatmap depicting the top 50 genes that are differentially expressed in *P. melaninogenica* upon its co-culture with the CF polymicrobial community model, composed of *P. aeruginosa*, *S. aureus*, *S. sanguinis*, and *P. melaninogenica*, in either mucin-containing ASM (+Mucin; this condition was referred to as “Mix” in the original publication (32)) or ASM lacking mucin (-Mucin; this condition was referred to as “Mix_base” in the original publication (32)). **B.** The *P. melaninogenica* genes highlighted in red are significantly downregulated in the absence of mucin, and include genes involved in metabolism of acetate and succinate, as well as pathways associated with the TCA cycle, pentose phosphate and serine metabolism.

Upon performing a gene-list enrichment analysis (33) of the most differentially expressed genes in the presence of mucin (**Figure 6A**, red blocks at top), we found that a number of those genes belonged to pathways involved in the catabolism of acetate and succinate (**Figure 6B**), the products of the *P. aeruginosa* malonate and propionate metabolism, respectively, identified above. Additionally, we noted the increased expression of genes encoding proteins associated with the TCA cycle, the pentose phosphate pathway, and serine catabolism (**Figure 6B**), indicating a general uptick in carbon metabolism by *P. melaninogenica* in the presentence of mucin when part of the polymicrobial community.

In addition to the overall increase in metabolism, we identified a particular CAZyme (34) that was significantly increased in expression in *P. melaninogenica* in the presence of mucin. The glycoside hydrolase HMPREF0659_A5155 belongs to the glycoside hydrolase 77 (GH77) family (34) and is predicted to be a putative 4-α-glucanotransferase which can transfer a segment of a 1,4-α-D-glucan onto the 4-position of an acceptor molecule (35, 36). To investigate the transcription of this enzyme in *P. melaninogenica*, we grew the organism anaerobically in co-culture with *P. aeruginosa* in mucin-containing ASM and measured the fold change of the expression of the gene via over time RT-qPCR. It was evident that the gene was not expressed in the early stages of the co-culture before displaying a ∼5-fold increase at the 6-hr time point, then returning to around baseline by the end of the co-culture (**Figure 7**). This observation coincides with the colony counts of the previous time course assay where *P. melaninogenica* appeared to require a 6-hr lag period before growing in co-culture with *P. aeruginosa* (**Figure 1B**).

**Figure 7.**
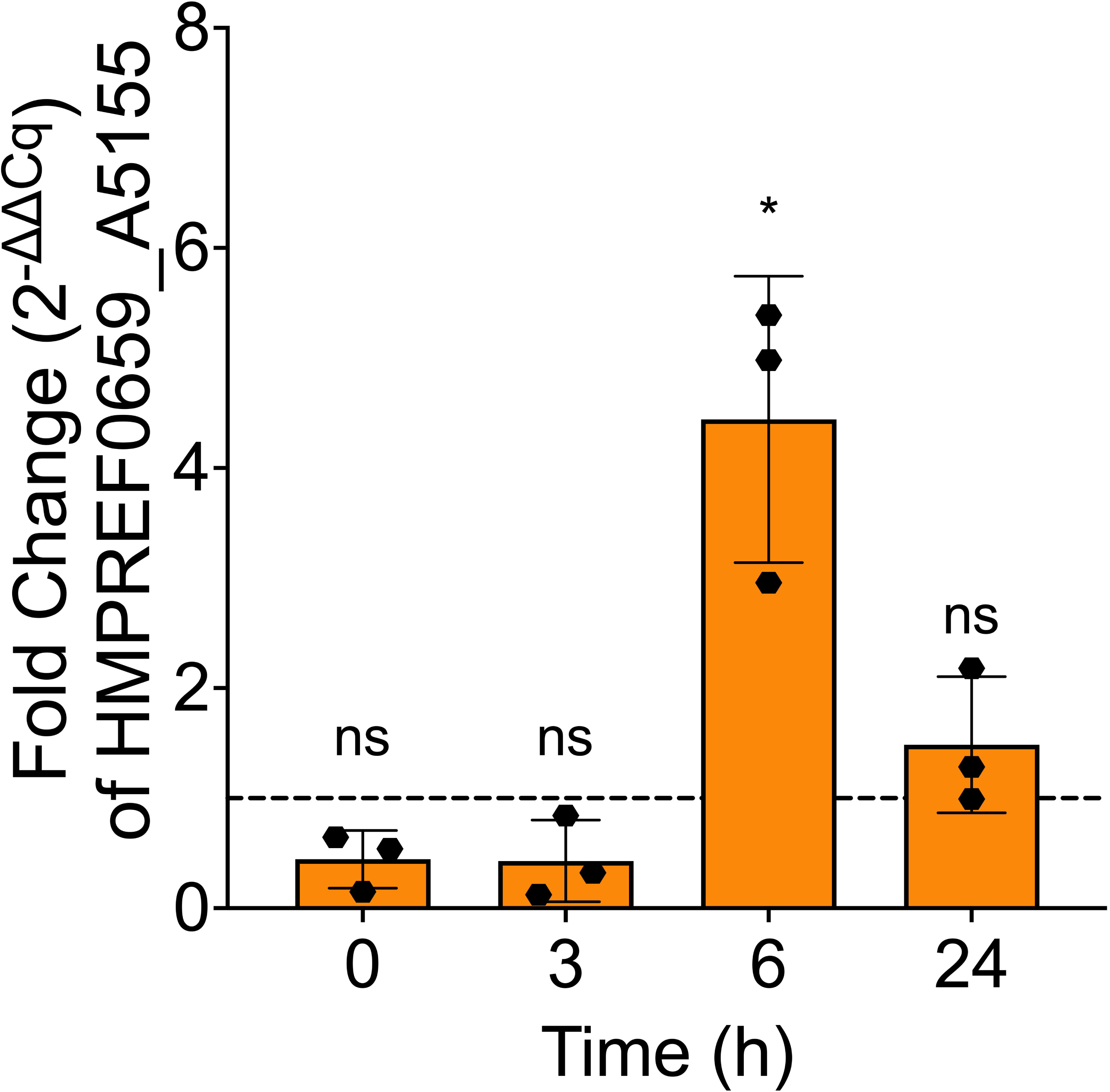
The *P. melaninogenica* CAZyme HMPREF0659_A5155 is upregulated in the presence of mucin during co-culture with *P. aeruginosa*. There is an approximate 4.5x increase in the fold change in gene expression of HMPREF0659_A5155 when *P. melaninogenica* is co-cultured with *P. aeruginosa* in mucin-containing ASM at the 6-hr time point, suggesting active mucin metabolism during co-culture. Statistical significance was calculated using two-tailed student’s t-test, * p < 0.05.

### Mucin glycans and amino acids are sufficient to support the growth of *P. melaninogenica* in co-culture with *P. aeruginosa*

The experiments above indicated that in the context of ASM, added mucin is required for *P. aeruginosa* to support the growth of *P. melaninogenica* and that growth in mucin induces the genes required for catabolism of succinate and acetate. Together, these data indicate that it is the catabolism of mucin that drives the interactions between these microbes. However, the complex and hard-to-characterize nature of mucin complicates our interpretations, prompting us to further simplify the medium to better understand the role of mucin *P. melaninogenica-P. aeruginosa* interactions. Considering the glycoprotein nature of mucins (37), we investigated the capacity of mucin components (glycans and amino acids) to support the growth of *P. melaninogenica* in co-culture with *P. aeruginosa* in a minimal salts medium with glycerol and nitrate under anoxic growth conditions. The selected mucin glycan sugars were N-acetylgalactosamine, N-acetyl-glucosamine, galactose, and fucose (38), while the amino acid component of mucin (37) was represented by casamino acids (CAAs). The co-culture of *P. melaninogenica* with *P. aeruginosa* in minimal medium containing mucin allowed for the growth of the former to ∼10^5^ CFU/mL (**Figure 8)**. Interestingly, *P. melaninogenica* was also recoverable, albeit to a significantly lesser amount (3.16×10^3^ CFU/mL), when co-cultured with *P. aeruginosa* in minimal medium containing both mucin glycans and CAAs, but not either component individually (**Figure 8**). *P. melaninogenica* was undetectable in all monoculture conditions (data not shown). Here, *P. aeruginosa* was recovered from mono- and co-cultures regardless of whether mucin or its components were present in the minimal medium (**Figure S8**). However, *P. aeruginosa* grew to a significantly lesser extent with mucin components compared to mucin in both mono- and co-culture conditions (**Figure S8**). Taken together, these data show that a simplified minimal salts medium with mucin components can largely recapitulate the observed *P. melaninogenica-P. aeruginosa* interaction in ASM + mucin. Furthermore, these data demonstrate that mucin, or carbon compounds derived from mucin, are a major mediator in the interaction between these organisms.

**Figure 8.**
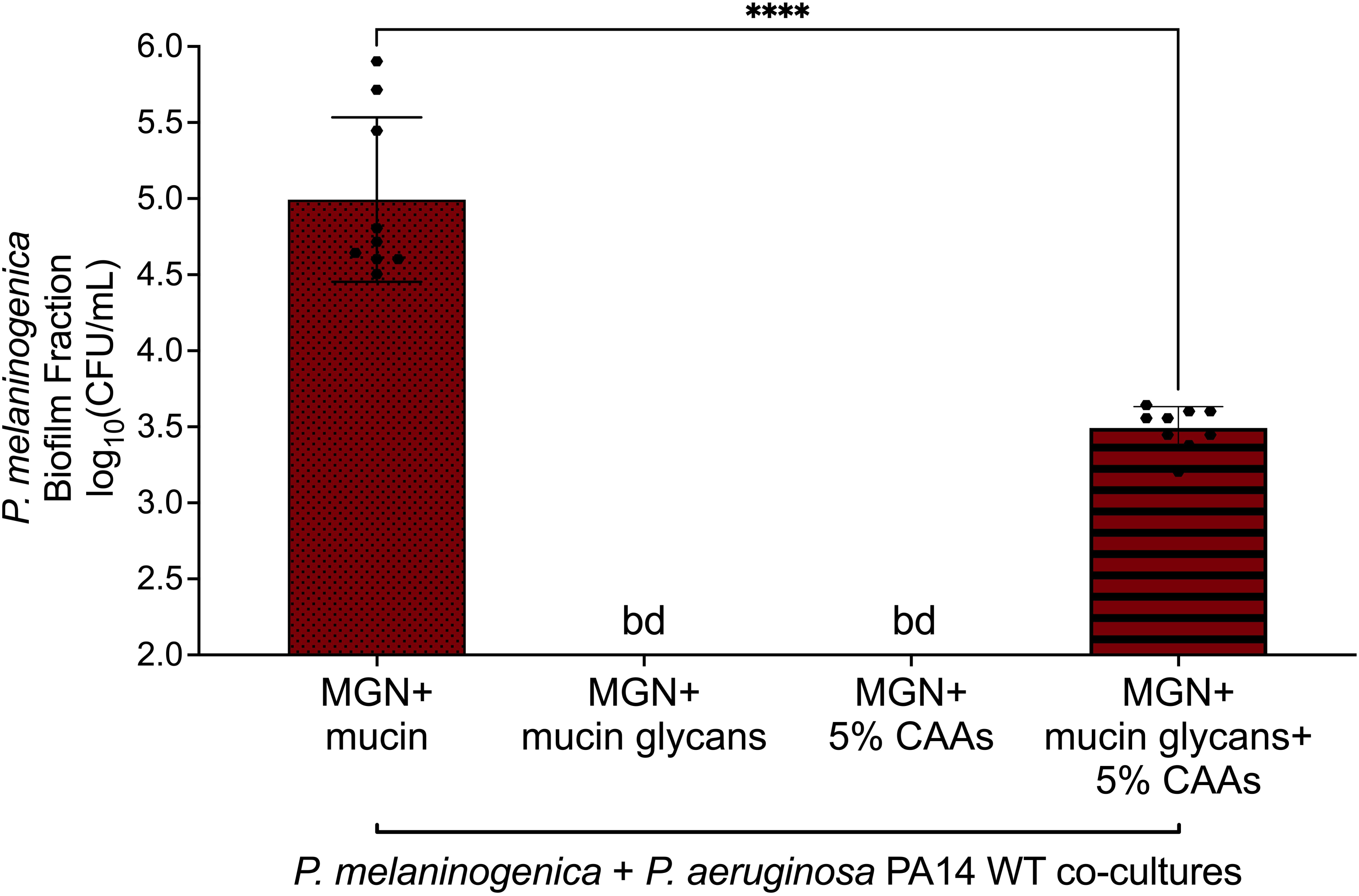
*P. melaninogenica* requires both the sugar and amino acid components of mucin to support its growth in co-culture with *P. aeruginosa*. The growth of *P. melaninogenica* in co-culture with *P. aeruginosa* can be partially supported with a medium containing the mucin components mucin glycans (fucose, galactose, N-acetylglucosamine, and N-acetylgalactosamine) and amino acids (casamino acids, CAA). Statistical significance was calculated using two-tailed student’s t-test, **** p < 0.0001.

## Discussion

In this study, we leveraged an existing CF airway polymicrobial community model, consisting of *P. aeruginosa, S. aureus, S. sanguinis,* and *P. melaninogenica* (11) to investigate the mechanisms that govern the ability of *P. melaninogenica* to grow in co-culture with the other members of the CF polymicrobial community, but not as a monoculture in mucin-containing ASM. By co-culturing *P. melaninogenica* in different combinations with other members, we identified *P. aeruginosa* as the main supporter of the survival of *P. melaninogenica* in the polymicrobial community. The inclusion of *P. aeruginosa* in the culture consistently improved the recovery of *P. melaninogenica* irrespective of the presence of *S. aureus* and *S. sanguinis*. We then investigated the temporal aspect of the interaction between *P. aeruginosa* and *P. melaninogenica*. The growth of *P. aeruginosa* in co-culture did not depend on the presence of *P. melaninogenica*; however, *P. melaninogenica* could not survive monoculture beyond 6 hours and required this first 6 hours for what appears to be a lag phase before growing in co-culture (**Figure 1B**).

To further elucidate the mechanism that governs the interaction between these organisms, we screened the *P. aeruginosa* PA14 non-redundant transposon mutant library (15) in search of a mutants that were incapable of supporting the growth of *P. melaninogenica* to the same extent as wild type *P. aeruginosa* PA14 in co-culture. We identified carbon and amino acid metabolism in *P. aeruginosa* PA14 as key pathways implicated in the interaction between these organisms (**Table S1**). Since previous metabolic modeling of the CF-relevant polymicrobial community predicted that organic acids were predicted to be cross-fed between members of the model community (8, 11), we experimentally pursued the malonate and propionate metabolism pathways that were identified by the screen (**Figure 2B**). To that end, we co-cultured *P. melaninogenica* with *P. aeruginosa* PA14 deletion mutants of the *mdc* and *prp* genes, rendering them incapable of converting malonate to acetate and propionate to succinate, respectively. The viability of *P. melaninogenica* was significantly reduced by ∼10 fold when anoxically co-cultured in mucin-containing ASM with any of *P. aeruginosa* PA14 Δ*mdcA,* Δ*mdcC*, Δ*mdcE,* Δ*prpB,* and Δ*mdcC*Δ*prpB* mutants compared to WT *P. aeruginosa* PA14 (**Figure 2C-D**). Those observations indicated that the disruption of acetate and succinate production in *P. aeruginosa* can significantly handicap the growth of *P. melaninogenica* in co-culture under CF-like conditions. Thus, acetate and succinate, as well as their respective metabolic precursors, malonate and propionate, seem to be mediators that are shared between organisms to help *P. melaninogenica* survive the CF polymicrobial environment. Additionally, we demonstrated that other metabolic pathways in *P. aeruginosa* that result in the formation of acetate and succinate (**Figure 3A**), namely those that involve *pauA*, *sucDC*, and *sdhBADC*, can also impact the interaction between *P. aeruginosa* and *P. melaninogenica*. We co-cultured *P. melaninogenica* with *P. aeruginosa* PA14 Δ*pauA*, Δ*sucDC*, Δ*prpB*Δ*mdcC*, Δ*sucDC*Δ*prpB*, Δ*sucDC*Δ*mdcC*, Δ*mdcC*Δ*pauA*, Δ*sucDC*Δ*prpB*Δ*mdcC*, Δ*prpB*Δ*mdcC*Δ*pauA*, and Δ*prpB*Δ*mdcC*Δ*sdhBADC* mutants (**Figure 3B**), and observed that *P. melaninogenica* exhibits reduced viability upon its co-culture with those *P. aeruginosa* PA14 mutants in mucin-containing ASM.

Interestingly, supplementing acetate or succinate at 6 hours into the co-culture of *P. melaninogenica* with either PA14 Δ*mdcC,* Δ*prpB*, Δ*pauA*, Δ*prpB*Δ*mdcC*, Δ*mdcC*Δ*pauA*, Δ*sucDC*Δ*prpB*Δ*mdcC*, Δ*prpB*Δ*mdcC*Δ*pauA*, or Δ*prpB*Δ*mdcC*Δ*sdhBADC*, but not Δ*sucDC,* Δ*sucDC*Δ*prpB*, or Δ*sucDC*Δ*mdcC,* restored the *P. melaninogenica* viable counts to those observed when it was co-cultured with WT *P. aeruginosa* PA14 (**Figures 2C-D & 3B-C**). This 6-hr time point aligns with the time course co-culture assay results, indicating that the interaction between *P. melaninogenica* and *P. aeruginosa* requires some time for *P. melaninogenica* to adapt, and for certain metabolites to be produced by the organisms before they are cross-fed to support its growth. We do not understand why supplementing the Δ*sucDC,* Δ*sucDC*Δ*prpB*, or Δ*sucDC*Δ*mdcC* with succinate did not rescue growth, but speculate that alterations in succinyl-CoA production might be having unanticipated effects on cell physiology.

We were able to establish that supernatants from co-cultures of *P. melaninogenica*-*P. aeruginosa*, but not either organism alone, could support the growth of *P. melaninogenica* in monoculture (**Figure 1C**), indicating that need for the close interaction of both organisms in this cross-feeding mechanism. Additionally, we were able to measure increased concentrations of acetate and succinate in the *P. melaninogenica*-*P. aeruginosa* co-culture supernatant (**Figure 4**), further supporting the metabolite cross-feeding model. The detection of acetate and succinate at higher concentrations in the co-cultures of our CF-like *in vitro* model also has physiological relevance. A previous study reported that the median acetate concentration was found to be more than double in the exhaled breath of 58 pwCF at 178 parts-per-billion by volume (ppbv) compared to healthy controls at 80 ppbv (39). Likewise, succinate was determined to be 10-fold higher in the sputa of 23 pwCF compared to 19 healthy controls (40). And thirdly, the concentrations of short-chain fatty acids (SCFAs) in the sputa of 9 pwCF was previously reported to be between 0.82 - 4.06 mM (41), which is near the range of the concentrations we detected in our studies here.

However, the cross-feeding of acetate and succinate does not fully explain the mechanism underlying the interaction between *P. melaninogenica* and *P. aeruginosa* since the disruption of their metabolic pathways, either individually or in conjunction, did not result in the complete loss of *P. melaninogenica* detection in co-cultures as observed with monocultures. Also, the addition of both acetate and succinate to *P. melaninogenica* monocultures did not rescue the growth of this anaerobe (**Figure S2C**), indicating the involvement of additional metabolic pathways in the interaction process. Therefore, we co-cultured *P. melaninogenica* with *P. aeruginosa* PA14 carbon catabolite repression mutants to evaluate the effect of disrupting broader metabolic pathway regulation on the interaction between the organisms. We observed that *P. aeruginosa* PA14 Δ*cbrA,* Δ*cbrB* and Δ*crc* mutants supported the growth of *P. melaninogenica* to a significantly lesser extent than WT *P. aeruginosa* PA14 (**Figure S5**). Such finding highlights the complexity of the mechanisms governing the interaction between *P. melaninogenica* and *P. aeruginosa* since CCR is involved in regulating the uptake and metabolism of different carbon sources, amino acids, lipids, and nucleic acids, as well as phenazines biosynthesis and the PQS system (30, 42). Interestingly, some CCR targets include *ascA* and *prpC* (30, 43), which are involved in propionate metabolism, *hutU* (44), which is involved in histidine metabolism, and *dctA*, which is a C_4_-dicarboxylate transporter of succinate, fumarate, and malate (30, 31), highlighting the importance of the ability of *P. aeruginosa* to transport organic acids when cross-feeding *P. melaninogenica*.

We were also able to demonstrate that *P. aeruginosa* does not utilize the mucin found in ASM since its growth as a monoculture was not impacted by the presence of mucin (**Figure 5A**). These data align with previously published observations (27). Interestingly, that same publication also demonstrated that a CF-derived consortium of anaerobic bacteria, which includes *P. melaninogenica*, was able to ferment mucin into the SCFAs acetate, propionate, and lactate (27).

By coupling these observation with experimental data indicating the dependency of *P. melaninogenica* on mucin in co-culture with *P. aeruginosa* (**Figure 5B-C**, **Figure 6**) and prior transcriptomic data showing the reliance of *P. melaninogenica* on mucin in ASM (32), along with RT-qPCR data that indicate the capability of *P. melaninogenica* to induce the expression of CAZymes implicated in mucin catabolism (**Figure 7**), as well as our data suggesting that only the *P. aeruginosa*-*P. melaninogenica* co-culture supernatants support the growth of *P. melaninogenica* monocultures **(Figure 1C)**, we propose a model of interaction between *P. aeruginosa* and *P. melaninogenica* in our CF-relevant community that relies on a two-way cross-feeding mechanism. That is, *P. melaninogenica* first ferments mucin to malonate and propionate (likely during the first 6 hours of the co-culture) before *P. aeruginosa* metabolizes malonate and propionate into acetate and succinate, respectively, and cross feeds these metabolites to *P. melaninogenica* to allow its growth in a CF-like environment (**Figure 9**).

**Figure 9.**
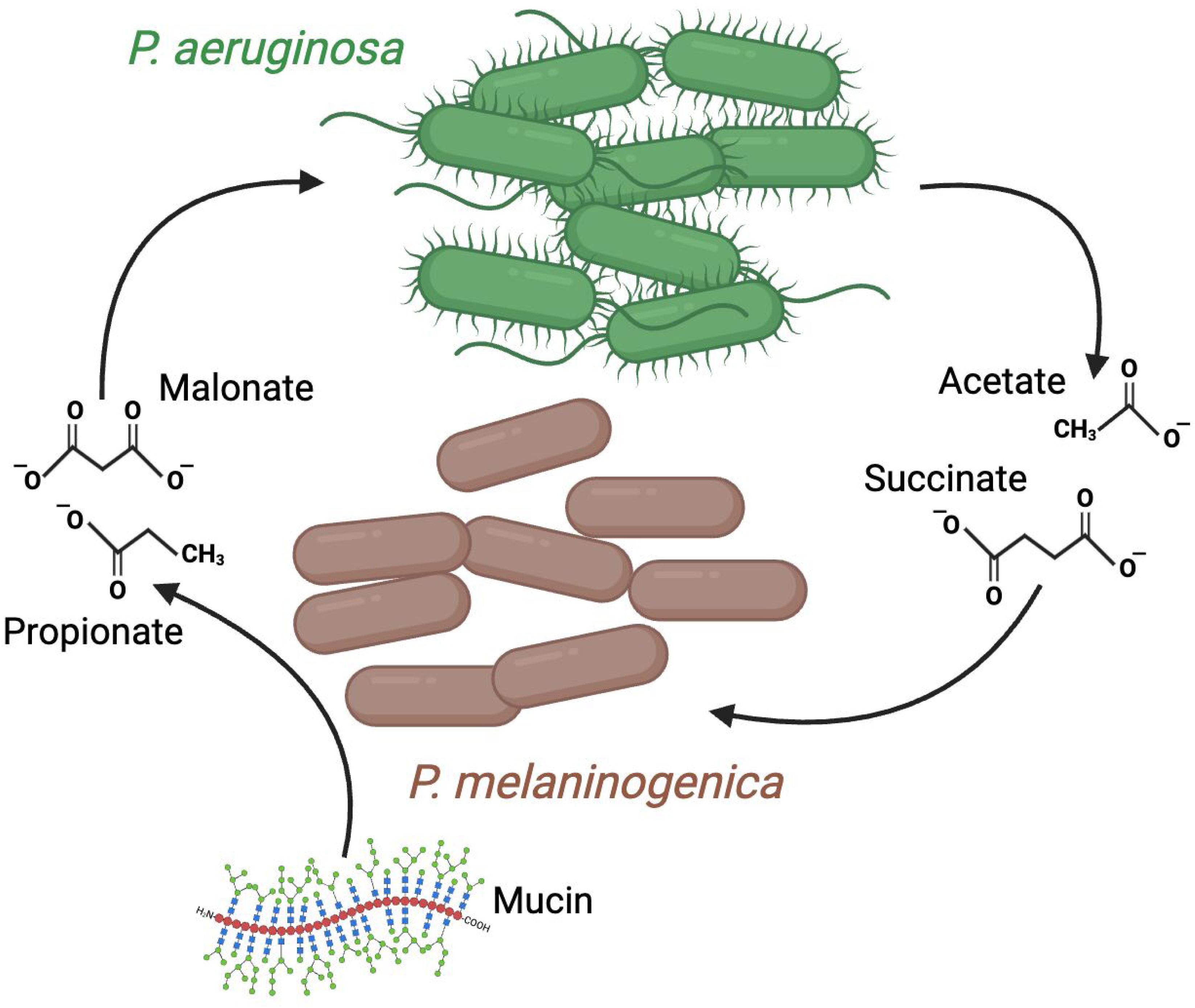
A *P. aeruginosa*-*P. melaninogenica* metabolic cross-feeding model. Based on the data presented here, we propose the following model. *P. melaninogenica* initiates the interaction with *P. aeruginosa* by fermenting mucin to malonate and propionate, which are then metabolized by *P. aeruginosa* into acetate and succinate, respectively. Acetate and succinate then serve as growth substrates for *P. melaninogenica* to assist in its growth in a CF lung-like environment. Figure designed using BioRender.

## Materials & Methods

### Bacterial strains and culture conditions

*P. aeruginosa* PA14 (45), *S. aureus* Newman (46), *S. sanguinis* SK36 (47), and *P. melaninogenica* ATCC 25845 (48) were used in this study and cultured in accordance with previously described methods (11). Briefly, *P. aeruginosa* and *S. aureus* were grown overnight in LB (lysogeny broth) at 37°C with shaking. *S. sanguinis* was grown overnight in Todd-Hewitt broth with 0.5% yeast extract at 37°C with 5% CO_2_. *P. melaninogenica* was grown anoxically overnight at 37°C in modified tryptic soy broth yeast extract (TSBYE), composed of tryptic soy broth (TSB) with 0.5% yeast extract, 5 µg/mL hemin, 500 µg/mL L-cysteine, and 1 µg/mL menadione. The list of strains used in this study can be found in **Table S2**.

### Bacterial co-culture assays

All co-cultures assays conducted in this study followed a procedure similar to what was previously published (11), with specific adjustments made to suit each experiment as detailed in the text. Generally, assays were performed in 96-well plates, overnight liquid cultures of the test bacterial strains were collected, pelleted and washed twice with 1X PBS, except for *P. melaninogenica* and *S. sanguinis*, which were washed once. Afterwards, the optical density (OD_600_) of the cultures were normalized to 0.05 in either mucin-containing ASM or minimal medium. The OD-normalized cultures were then either dispensed into the 96-well plate for a final OD_600_ = 0.01 or mixed together so that each member would have a final OD_600_ = 0.01, and then the mixture was dispensed into the 96-well plate. Plates were then enclosed in Thermo Fisher Scientific AnaeroPack^TM^ anaerobic boxes along with a BD GasPak^TM^ anaerobe sachet and incubated at 37°C for 24 hours. Following incubation, the planktonic fractions of the cultures were separated from the biofilm fraction in the 96-well plate, 50 μL of 1X PBS were added to each test well, and the biofilm was mechanically detached using a 96-pin replicator. The detached biofilm fractions then underwent 10-fold serial dilutions, and the entire dilution series were spotted onto selective media. CFUs were enumerated following overnight incubation, and the CFU per milliliter concentrations were determined.

The selective media used in the co-culture assays were as follows: *Pseudomonas* isolation agar (PIA) was used to recover *P. aeruginosa*, mannitol salt agar (MSA) was used to recover *S. aureus*, *Streptococcus* selective agar (SSA), made of blood agar supplemented with 10 μg/mL polymyxin B and 10 μg/mL oxolinic acid, was used to recover *S. sanguinis*, and *Prevotella* selective agar (PSA), composed of blood agar supplemented with 5 µg/mL hemin, 500 µg/mL L-cysteine, 1 µg/mL menadione, 100 µg/mL kanamycin, 7.5 µg/mL vancomycin, and 5 µg/mL polymyxin B, was used to recover *P. melaninogenica*. Cultures were incubated for 24 hours unless otherwise noted.

When co-culture supplementation experiments were performed with acetate and succinate, 100 mM stock solutions of sodium acetate and sodium succinate were prepared then diluted in mucin-containing ASM before being introduced into the cultures at final concentrations of 4.5 mM and 2.8 mM, respectively, at the 6-hr time point.

### Time course co-culture assays

The time course co-culture assays used in this study relied on the same experimental procedure as the endpoint co-culture assays described above; however, multiple plates in anaerobic boxes were inoculated in parallel, starting from the same overnight cultures, with each anaerobic box corresponding to a timepoint at which the cultures were processed as described above.

### P. aeruginosa-P. melaninogenica co-culture genetic screen

The transposon mutagenesis screen utilized the *P. aeruginosa* PA14 non-redundant transposon (Tn) mutant library (15) and was adapted from a previously described procedure (49) with modifications to accommodate *P. melaninogenica*. On the first day, the PA14 Tn library mutants were transferred into a 96-plate containing 100 µL of LB broth and incubated overnight at 37°C. In parallel, modified TSBYE liquid cultures of *P. melaninogenica* were started and incubated anoxically overnight at 37°C. On the second day, the *P. melaninogenica* cultures were OD-adjusted in mucin-contained ASM to an OD_600_ = 0.01, then dispensed into a 96-well plate. To that same plate, the PA14 Tn library grown overnight in LB was transferred using a 96-pin replicator. Thus, each well contained WT *P. melaninogenica* and a Tn mutant of *P. aeruginosa* PA14. The co-cultures were then incubated anoxically for 24 hours at 37°C. Following incubation, the planktonic fractions of the co-cultures were separated from the biofilm fractions, and 50 μL of 1X PBS were added to the co-culture plates. The biofilm fractions were then mechanically detached using a 96-pin replicator, and each co-culture plate was spotted onto PIA and PSA plates. The PIA plate was then incubated aerobically, and the PSA plate was incubated anoxically overnight at 37°C. Following incubation, *P. aeruginosa* Tn mutants that were recovered on the PIA plate but were unable to (or showed reduced ability to) support the growth of *P. melaninogenica* were identified. Candidate *P. aeruginosa* Tn mutants were individually retrieved from the transposon library and transferred to a separate 96-well plate to generate the primary candidate library. A second round of screening was performed using the same procedure, but with strains selected from the primary candidate library, followed by a third confirmatory test to generate the final list of *P. aeruginosa* PA14 Tn mutants that were unable to effectively support the growth of *P. melaninogenica* in co-culture (**Table S1**).

### *P. aeruginosa* PA14 gene deletions

The clean deletion mutants of *P. aeruginosa* PA14 *ΔmdcA*, *ΔmdcC,* and *ΔmdcE*, as well as the complementation mutant PA14 *ΔmdcC::mdcC* were acquired from the Dietrich Lab (20). The clean deletion mutants *P. aeruginosa* PA14 Δ*prpB*, Δ*acsA*, and Δ*prpB*Δ*acsA* were acquired from the Hunter Lab (27). The *P. aeruginosa* PA14 *ΔprpBΔmdcC* mutant was generated in the PA14 *ΔmdcC* background via conjugation with *E. coli* S17-1 harboring the deletion vector pSMV8::*prpB*-KO provided by the Hunter Lab (27). The *P. aeruginosa* PA14 Δ*pauA* and Δ*sucDC* were generated using via conjugation with *E. coli* S17-1 harboring either pMQ30::*pauA*-KO, pEX18Gm::*sucDC*-KO, or pEX18Gm::*sdhBADC*-KO made in-house. The combination mutants were created by conjugating different *P. aeruginosa* PA14 mutants with *E. coli* S17-1 harboring the desired deletion vector. The clean deletion mutants *P. aeruginosa* PA14 *ΔcbrA*, *ΔcbrB,* and *Δcrc* were acquired from the Hogan Lab (29) along with PA14 Δ*mvfR*/*pqsR* and Δ*pqsH*. The clean deletion mutant *P. aeruginosa* PA14 Δ*phzA1*Δ*phzA2* was acquired from the Newman Lab (50). The list of strains used in this study can be found in **Table S2**.

### Metabolite quantification

Supernatants resulting from the *P. aeruginosa-P. melaninogenica* co-cultures grown in mucin-containing ASM were collected, centrifuged, sterilized through a 0.22 μm filter, and frozen at −80°C in 1.5 mL microcentrifuge tubes. The samples were then shipped to the Mass Spectrometry and Metabolomics Core (MSMC) at Michigan State University where they were analyzed via gas chromatography-mass spectrometry (GC-MS) using protocols MSU_MSMC_010 and MSU_MSMC_010a. The organic acid concentrations were calculated by normalizing their values to standards, then normalizing those to blank ASM as a baseline.

Supernatants resulting from the *P. aeruginosa-P. melaninogenica* co-cultures grown in MGN plus mucin were collected, centrifuged, filtered through a 3 μm filter, then sterilized through a 0.22 μm filter. Cell-free supernatants were then analyzed via high-pressure ion chromatography (HPIC) using a Dionex™ IonPac™ AS11-HC-4μm column. The organic acid concentrations were calculated by normalizing their values to standards.

### Quantitative reverse transcription polymerase chain reaction (RT-qPCR)

Two separate *P. aeruginosa-P. melaninogenica* co-culture conditions were established as described above. One condition utilized mucin-containing ASM while the other utilized Modified TSBYE as culture medium. At the 24-hr time point, the QIAGEN RNeasy Mini Kit was used in accordance with manufacturer instructions to extract total RNA from the cells of both co-culture conditions. RT-qPCR was run using the following *P. melaninogenica*-specific primers: BH_rt_Pm_A5155_F: 5’-TAGGGTCAGCCAAACGCAAT-3’ and BH_rt_Pm_A5155_R: 5’-TTACATCGTGGTGGTCCTGC-3’ to target the CAZyme HMPREF0659_A5155, and BH_rt_Pm_gyrA_F: 5’-TTACACCGGGTACGTCAAGC-3’ and BH_rt_Pm_gyrA_R: 5’-GACACCGTGAGGAACTCTGG-3’ to target *gyrA* as a reference gene. The Livak (2^-ΔΔCT^) method (51) was used to calculate fold change in gene expression with the expression of the CAZyme in the Modified TSBYE co-culture condition acting as the control.

### Statistical Analysis

Analysis was performed using GraphPad Prism 10. The mean values ± standard deviations (SDs) were plotted. Either ordinary one-way analysis of variance (ANOVA) or student’s t-test were performed to determine statistical significance, as indicated in the figure legends.

## Acknowledgements

This work was supported by the National Institutes of Health (R01AI155424) to G.A.O. We thank the Dietrich lab for the *P. aeruginosa* PA14 Δ*mdcA*, Δ*mdcC*, and Δ*mdcE* mutants, as well as the complementation strain PA14 Δ*mdcC::mdcC*, the Hunter lab for the *P. aeruginosa* PA14 Δ*prpB*, Δ*acsA*, and Δ*prpB*Δ*acsA* mutants as well as the pSMV8::*prpB*-KO deletion vector, the Hogan lab for the *P. aeruginosa* PA14 Δ*cbrA*, Δ*cbrB*, Δ*crc*, Δ*mvfR*/*pqsR*, and Δ*pqsH* mutants, and the Newman Lab for the *P. aeruginosa* PA14 Δ*phzA1*Δ*phzA2* mutant. We also thank Dr. Christopher A. Kesthely for his assistance in visualizing and analyzing the data presented in **Figure 6**.

